# Low-frequency hearing thresholds improve as high-frequency hearing sensitivity deteriorates between young adulthood and middle age in normally hearing people

**DOI:** 10.1101/2024.03.06.583598

**Authors:** Tatiana Izmaylova, Jaime A. Undurraga, Paul F. Sowman

## Abstract

Hearing sensitivity changes throughout a person’s lifetime. This work aimed to describe changes in pure-tone audiometric (PTA) thresholds that occur in the transition from young adulthood to middle age in 121 adults with normal or nearly normal hearing. Results showed that older people had worse high-frequency (4000-8000 Hz) thresholds and better low-frequency (125-500 Hz) thresholds than younger individuals, suggesting that hearing sensitivity in the low-frequency range may improve with age. The improvement of low-frequency thresholds may be part of a central compensation for age-related deterioration of high-frequency hearing sensitivity. Further investigation of age-related changes in low-frequency hearing sensitivity is needed to confirm our findings.

## 1. Introduction

Hearing sensitivity decreases gradually with age. While one starts losing the ability to hear the highest frequencies early in life (Burén et al., 1992; Hemmingsen et al., 2021; Wang et al., 2021), hearing sensitivity within the conventional audiometric range (125-8000 Hz) remains normal until about 40 years of age (Brant & Fozard, 1990; Flamme et al., 2011; Kim et al., 2010; Wasano et al., 2021). As normal hearing sensitivity at and below 8000 Hz is considered sufficient for adequate speech perception (French & Steinberg, 1947), few studies have focused on subclinical changes in conventional hearing thresholds that occur between young adulthood and middle age. Yet, recent research suggests that these subtle changes in hearing may have more significant real-life implications than believed earlier. For example, variations in normal hearing sensitivity have been found to predict performance on speech-in-noise and cognitive tests (Chern et al., 2022; Golub et al., 2020; Holmes & Griffiths, 2019). Therefore, a better understanding of age-related changes in normal hearing sensitivity is necessary for predicting hearing and cognitive health outcomes.

The aim of this work was to examine changes in pure-tone audiometric (PTA) thresholds that occurred between young adulthood and midlife in the normally hearing population. More specifically, we were interested in tracking how hearing sensitivity changed with age at each audiometrically tested frequency. Results of past audiometric studies show that, by middle age, the most noticeable decline occurs at 8000 and 4000 Hz, while lower frequencies remain unaffected (Corso, 1959; Flamme et al., 2011; Kim et al., 2010; Morrell et al., 1996; Wasano et al., 2021). Based on those findings, we hypothesised that there would be no age-related changes in the low- and middle-frequency PTA thresholds and a reduction in hearing sensitivity at the two highest frequencies.

## 2. Materials and methods

### 2.1 Participants

One hundred and twenty-one people (57 females: mean age = 26.8 years, age range = 18-47 years; 60 males: mean age = 26.4 years, age range = 18-47 years) participated in the study. Participants were recruited for a neurophysiological study, and their thresholds were measured as part of the qualifying assessment. Only hearing threshold data is reported here. Most participants below 22 years of age were university students who participated in the study for course credit. The remaining participants were paid volunteers. Participants were included if their pure-tone air-conduction hearing thresholds averaged between the ears were <= 25 dB HL at octave frequencies between 125-8000 Hz. Four of the 121 people tested did not meet the inclusion criteria as their 8000 Hz or 4000 Hz averaged PTA thresholds were above 25 dB HL. The excluded participants were 21, 22, 37, and 53 years old. The information on the age distribution of the included participants can be found in Figure 1. Of the 117 included participants, 107 had their PTA thresholds at 20 dB HL or lower (normal hearing), while 10 people had some PTA thresholds (mainly, at 8000 Hz) between 20 and 25 dB HL (nearly normal hearing). The study was approved by the Macquarie University Ethics Committee (Reference No: 5201956309964).

**Fig. 1.**
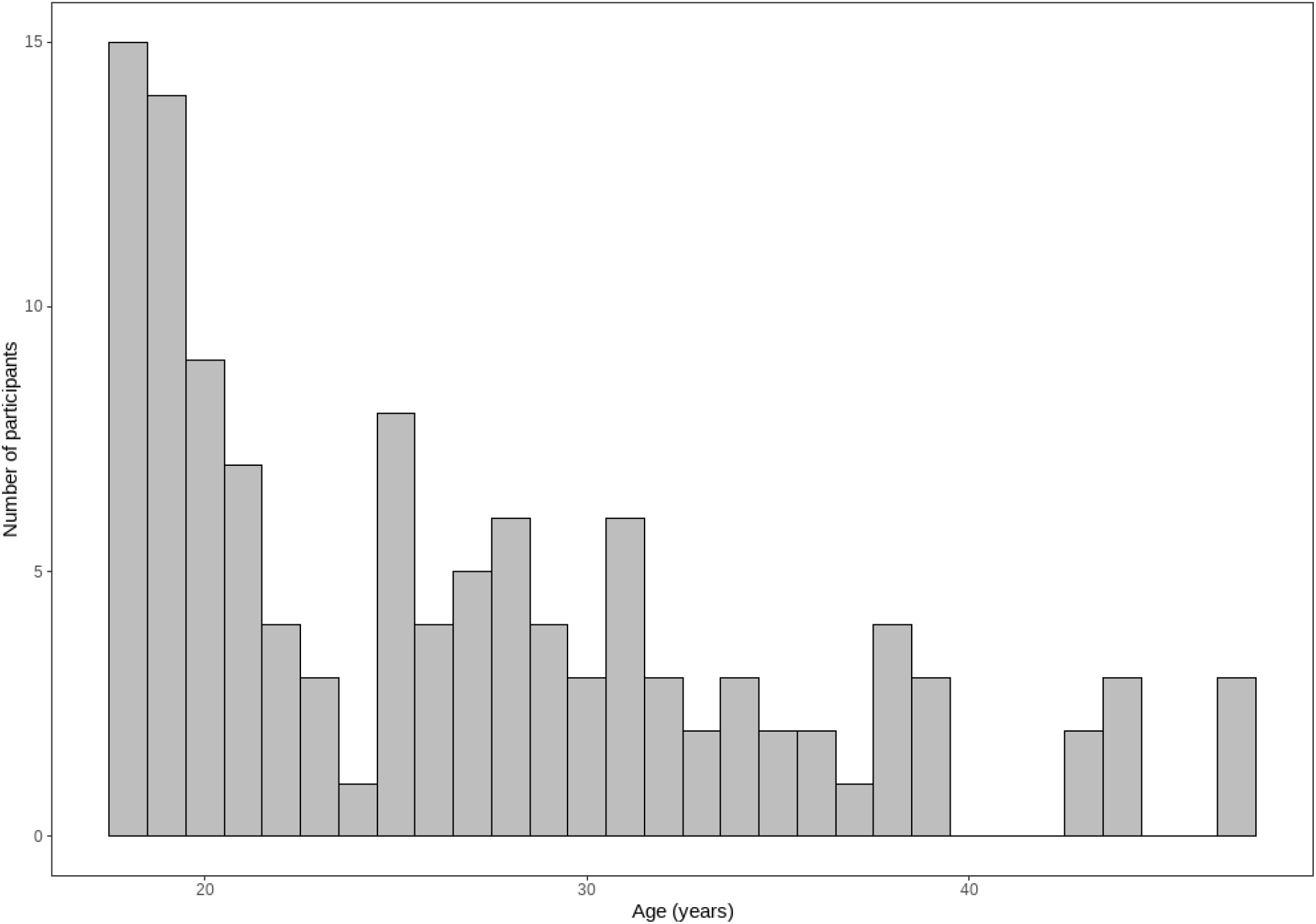
Participant age distribution.

### 2.2. Hearing assessment

The participant’s hearing examination was conducted in a soundproof booth using a calibrated clinical audiometer (Affinity 2.0, Interacoustics, Denmark) with a circumaural headset. Participants’ pure-tone air-conduction hearing thresholds were obtained for octave frequencies between 125 and 8000 Hz. We used a modified Hughson-Westlake method. The level was decreased in 10 dB steps starting from 40 dB HL. Stimulus tones were presented for 1-2 seconds, after which participants indicated positive responses with a button press. After a failure to respond, the level was increased in 5 dB steps until the stimulus was heard again. The threshold was the lowest level, where 2 out of 3 responses occurred.

### 2.3. Data analysis

All analyses were performed in R Statistical Software (v4.2.1; R Core Team 2022). Each participant’s PTA thresholds were averaged between the ears and analysed using a linear mixed-effects (LME) model (the lme4 package; v1.1-26; Bates et al., 2015). The advantage of the LME model is that it provides a more accurate estimation of effects due to modelling not only fixed but also random factors, which, in our case, would be participants’ individual differences in hearing sensitivity. The LME model included PTA thresholds as a dependent variable, whilst age (continuous) and frequency (7 levels: 125, 250, 500, 1000, 2000, 4000, 8000 Hz) were fixed factors. Participant ID was used as a random factor to account for between-subject variability. We measured the variance inflation factor (VIF) to control for possible collinearity across independent factors. The VIF did not exceed 4 for any factor, indicating low to moderate collinearity. Conditional, random, and marginal residuals of the LME model were checked by visual inspection. The Shapiro-Wilk test showed that the model’s residuals were not normally distributed. The data were log-transformed, which normalised the distribution of the residuals. Then, the model was refitted. Post-hoc analyses were adjusted for multiple comparisons using the ‘holm’ correction (Holm, 1979).

## 3. Results

Figure 2A shows age-related changes in the PTA thresholds at each tested frequency. As can be seen from the graph, the PTA thresholds appeared to increase (worsen) at the two highest frequencies (4000 and 8000 Hz) with age. Unexpectedly, the low-frequency thresholds (125 – 500 Hz) appeared to decrease (improve) between young adulthood and midlife.

**Fig. 2.**
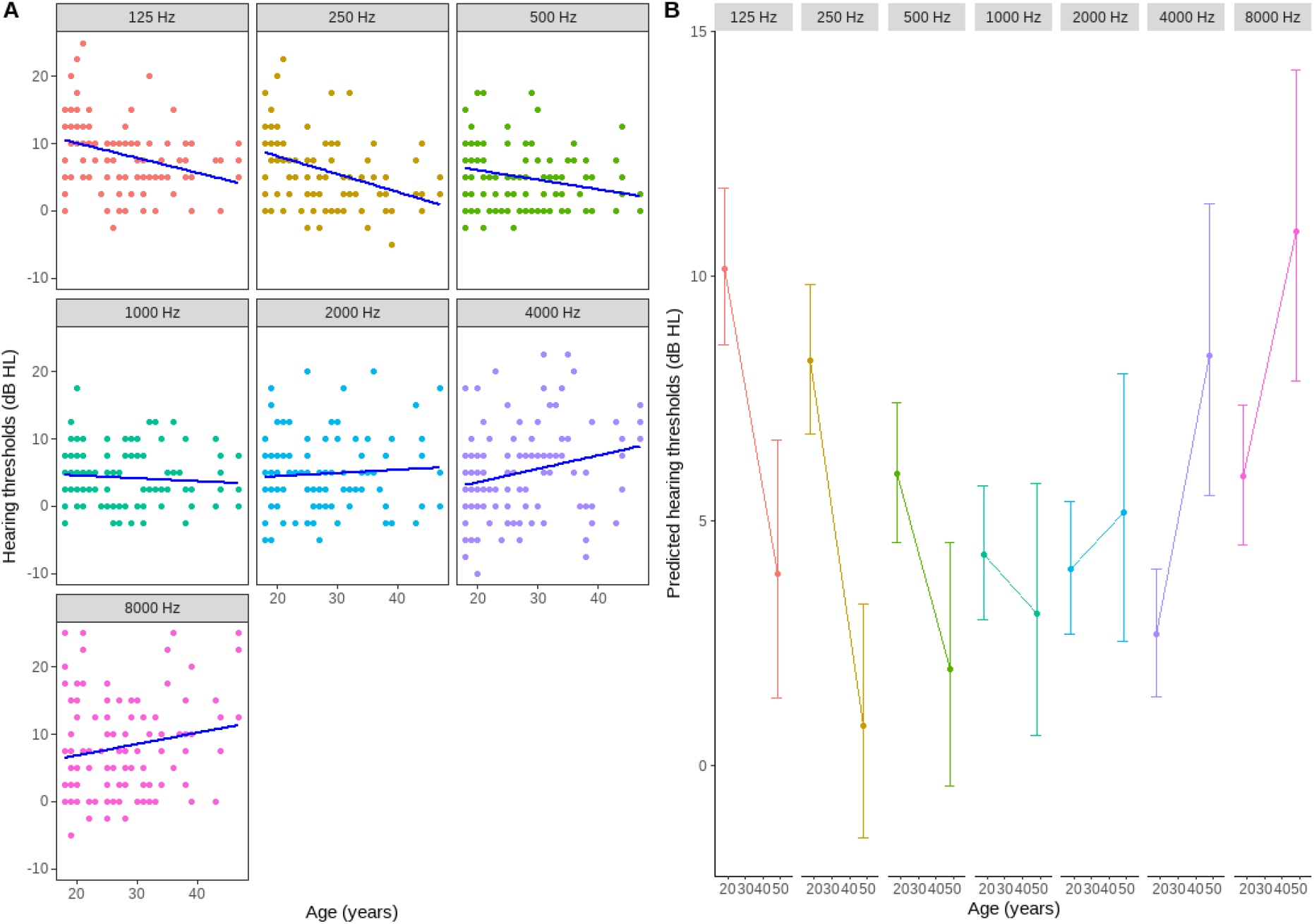
Age-related changes in PTA thresholds. (A) PTA thresholds at each tested frequency by age. The blue lines represent lines of linear regression. The frequency is indicated on the top of each panel and colour coded. (B) Model PTA predictions (via marginal means; Lenth, 2022) as a function of age for different frequencies (indicated on the top of each panel and colour coded). A significant frequency-by-age interaction was identified, whereby high-frequency PTA thresholds deteriorated and low-frequency thresholds improved between young adulthood and middle age. The error bars represent a 95% confidence interval.

This pattern of age-related changes was confirmed by the LME model. The results showed that there was a main effect of frequency [F(6, 690) = 13.29, p < .0001] and a frequency by age interaction [F(6, 690) = 10.79, p < .0001]. Post-hoc analysis revealed a significant age-related reduction in hearing sensitivity at 4000 Hz [t(562) = -3.09, p = .002] and 8000 Hz [t(562) = -2.51, p = .01]. Conversely, the PTA thresholds improved significantly at 125 Hz [t(562) = -3.25, p = .0012], 250 Hz [t(562) = --4.16, p < .0001], and 500 Hz [t(562) = -2.27, p = .02] between young adulthood and middle age. No age-related changes in the middle-frequency (1000-2000 Hz) thresholds were found. The model’s predictions for age-related changes in PTA thresholds by frequency are shown in Figure 2B.

## 4. Discussion

This work explored age-related changes in hearing sensitivity between young adulthood and middle age in normally-hearing adults. We hypothesised that there would be an age-related increase in PTA thresholds at 8000 and 4000 Hz, with no changes at other tested frequencies. The hypothesis was only partially supported. As expected, the high-frequency thresholds gradually declined with age. However, the low-frequency thresholds showed an age-related improvement which was not anticipated.

Enhancement of low-frequency hearing sensitivity has not been a topic of discussion in hearing research. To our knowledge, two studies by Morrell et al. (1996) and Corso (1959) mentioned a slight improvement of PTA thresholds (mainly in the low- and middle-frequency range) between young adulthood and middle age in females and males, respectively. However, no attempts were made to explain the unusual observations due to a descriptive focus of those studies. Thus, whether the age-related enhancement of hearing sensitivity reported in those studies and our results are incidental or part of a yet-to-be-confirmed trend remains unclear.

One very speculative explanation for the low-frequency enhancement around middle age is compensatory changes that may occur in the central auditory system following peripheral hearing loss in the high-frequency range. Some support for this theory comes from hearing loss studies in mice. Willott (1986) conducted multi-unit recordings to investigate changes in frequency representation by neurons in the inferior colliculus (IC) of mice genetically predisposed to early hearing loss. They discovered downward shifts of at least 1 octave in the best frequencies of neurons of the older mice in the regions usually processing high-frequency sounds. Remarkably, the mean response thresholds of the IC low-frequency neurons became significantly lower (better) as high-frequency thresholds declined. A similar shift to increased representation of lower frequencies was found in the auditory cortex and the dorsal cochlear nucleus of the predisposed to hearing loss mice (Bishop et al., 2022; Willott et al., 1991, 1993). The increased low-frequency sensitivity eventually disappeared following the progression of hearing loss. At the behavioural level, mice predisposed to hearing loss exhibited increasingly exaggerated acoustic startle responses to low-frequency pips as their hearing loss progressed. In contrast, normally hearing mice continued to respond strongly to mid- and high-frequency stimuli (Ison & Allen, 2003). Similarly, low-frequency tones played before startle-evoking stimuli became more efficient at attenuating startle responses with age in mice predisposed to hearing loss, while normally hearing mice did not experience changes in startle modification (Willott et al., 1994). Overall, the findings of Willot and colleagues demonstrate that temporary age-related improvement of sensitivity to low frequencies is possible at cortical and subcortical levels after the onset of high-frequency peripheral hearing loss. It is plausible that compensatory changes in the central auditory system could temporarily reduce low-frequency hearing thresholds. This possibility aligns with our results and is further supported by studies on age-related changes in the acoustic startle reflex in mice.

The idea that neuroplastic changes in the auditory system following hearing loss may benefit perception is not new. Moore and Vinay (2009) reported that people with cochlear lesions (termed “dead regions”) showed enhanced frequency discrimination abilities for frequencies located just below the lesions. They were also better at amplitude-modulation detection and consonant discrimination when those stimuli were restricted to the low-frequency range. Moore and Vinay (2009) concluded that the improved performance in people with cochlear dead regions was due to the remapping of the auditory cortex, whereby the lower frequencies adjacent to the lesion became overrepresented. While our participants had normal hearing in the conventional audiometric range, they may have experienced age-related hearing loss at high and extended frequencies. Hearing sensitivity in the extended high-frequency range begins to decline during school years (Burén et al., 1992; Hemmingsen et al., 2021) and by middle age, a significant decrement is accrued (Rodríguez Valiente et al., 2014). The gradual improvement of the low-frequency hearing thresholds between young adulthood and middle age may be related to the remapping of the auditory cortex initiated by the reduction of hearing sensitivity at high frequencies.

If we consider the possibility of compensatory changes in the central auditory system, a logical question would be why enhanced low-frequency hearing sensitivity measured as pure-tone air- or bone-conduction hearing thresholds has not been more frequently reported in the hearing research. One possible explanation is that the compensatory changes in the central auditory system may be affected by individual differences in the trajectory of hearing loss. As Willott and colleagues noted, only the mice genetically predisposed to hearing loss showed a significant shift to lower frequencies across the auditory system. At the same time, no such changes were found in the mice without the predisposition to hearing impairment, even when their hearing sensitivity started deteriorating with age. Similarly, humans may have different degrees of predisposition to hearing loss (Barrenäs & Lindgren, 1990; Varghese & Kottaramveettil, 2019), which may introduce variability in the onset of the compensatory changes. Noise exposure is another factor that may contribute to an earlier onset of hearing loss that may alter the timeline of auditory compensation. Therefore, for the compensatory changes to be noticeable, one may need a relatively homogeneous population sample to minimise the effect of the genetic and environmental factors. In our study, participants were predominantly White persons, which ensured that potential race-related differences in hearing sensitivity (e.g., due to variations in melanin concentrations in the inner ear known for its protective effect on hearing; Agrawal et al., 2008; Helzner et al., 2005; Lin et al., 2012) were minimised. Additionally, all our participants were university-educated, which made them less likely candidates for exposure to occupational noise, thus reducing variability in the onset of hearing loss and resulting compensatory changes. So, there is a possibility that the racial and educational homogeneity of our sample may have contributed to identifying the subtle age-related enhancement of low-frequency hearing sensitivity that would be difficult to detect in a more heterogeneous population.

An alternative explanation to our findings could be that the difference observed between the younger and older people was incidental due to an unknown sample selection error. Indeed, the main limitation of this study was its cross-sectional nature, with the data coming from different people aged between 18 years and midlife. The youngest participants were undergraduate students who participated in the study for course credit. Other participants were paid volunteers partly recruited from graduate students and partly from social media. There is a possibility that the older participants were more motivated to show good results on the hearing test, thus creating a bias in the data. However, this scenario is unlikely as such bias would probably not be frequency specific.

It should be noted that the observed results were not due to a higher exclusion rate among the older participants than younger ones. Only four people from the entire sample did not meet the inclusion criteria, two of whom were in their 20s, one in their 30s, and one in their early 50s. This fact eliminates the possibility that the inclusion criteria favoured the younger participants, creating a selection bias for the older ones. At the same time, an unequal distribution of participants across the age groups, resulting in the broadest variability of hearing sensitivity in the youngest participants, could contribute to the study’s outcome. As it is unclear whether these or any other factors may have inadvertently affected our results or whether our findings reflected the neural compensation for the reduced hearing sensitivity, more research is needed. Future studies would focus on analysing large cross-sectional datasets consisting of participants with the same socio-economic backgrounds and longitudinal data.

## Supporting information

Hearing Thresholds Data File

## Authorship contribution statement

Tatiana Izmaylova: Conceptualisation, Methodology, Investigation, Formal analysis, Visualisation, Writing – original draft preparation, Writing – review & editing. Jaime A. Undurraga: Methodology, Resources, Writing – review & editing, Supervision. Paul F. Sowman: Writing – review & editing, Supervision.

## Acknowledgements

This research was supported by an Australian Government Research Training Program (RTP) Scholarship.

